# External Evaluation of Multi-Omics Prognostic Models Across TCGA-BRCA and METABRIC: Transportability, Stability, and Incremental Performance

**DOI:** 10.64898/2026.09.18.752709

**Authors:** Elena Krikun, Abedalrhman Alkhateeb

**Author notes:** Corresponding author. *Email addresses:* (Elena Krikun). These authors contributed equally to this work. (Abedalrhman Alkhateeb).

## Abstract

Cross-cohort evaluation of multi-omics prognostic models can fail because molecular features are not assayable, fitted models do not transport, feature selection is unstable, or molecular data add little beyond clinical predictors. We used a dual-track evaluation design with TCGA-BRCA as the source cohort and METABRIC as the independent target cohort. Track A tested outcome-blind transport of fixed historical RNA and copy-number models without target-cohort refitting, whereas Track B reconstructed the dependency-aware selection procedure within METABRIC using leakage-controlled repeated cross-validation. Literal transport provided little or negative incremental overall-survival discrimination: RNA reduced Harrell’s C-index relative to clinical prediction (Δ*C* = *−*0.0136), while copy number was essentially neutral. Track B showed no reliable overall-survival gain for RNA, copy number, or mutation; methylation and the reconstructed multimodal model performed worse than matched clinical comparators. In a protocol-locked post-hoc extension, the same fixed features were retained but model parameters were re-estimated within METABRIC. Incremental C-index increased by 0.0162 for RNA, 0.0091 for copy number, and 0.0201 for the combined panel. Random-panel benchmarks showed that the RNA gain was common among alternative panels, whereas the historical copy-number panel ranked above all 200 panels sampled from its recovered assayable candidate space. The recurrence-free-survival sensitivity analysis showed a positive RNA contrast (Δ*C* = 0.0146). These results separate failure of fixed-model transport from loss of information in the underlying feature set and show that successful local redevelopment does not by itself establish feature-set specificity.

## 1. Introduction

Molecular profiling has become an important component of breast-cancer prognostic research. Gene-expression signatures can stratify patients by outcome, but strong performance in a development cohort does not guarantee that the same features or fitted model will remain useful in an independent population [1]. This distinction is particularly important in high-dimensional molecular data, where many correlated feature sets can capture overlapping disease structure. Indeed, randomly constructed gene-expression signatures have also been associated with breast-cancer outcome [2].

Cross-cohort evaluation becomes more difficult when several molecular modalities are involved. TCGA-BRCA and METABRIC are both extensively characterized breast-cancer cohorts, but they differ in assay platforms, feature coverage, identifiers, missingness, clinical composition, and follow-up [3, 4, 5]. Consequently, poor performance in an external cohort can have several explanations. A source feature may not be measurable on the target platform; a measurable feature set may retain information while its fitted coefficients fail to transport; a feature-selection procedure may identify unstable representations; or molecular measurements may simply add little prognostic information beyond established clinical variables.

These possibilities should not be treated as a single external-validation outcome. Large multi-omics survival benchmarks have shown that adding high-dimensional molecular data does not consistently improve prediction over clinical models across cancer datasets [6]. More recent standardized benchmarking has reached a similar conclusion: performance depends strongly on the cohort, available modalities, clinical covariates, and evaluation design [7]. A useful cross-cohort analysis therefore needs to identify *why* a molecular model does or does not reproduce, rather than reporting external performance alone.

In our previous work, we developed a dependency-aware Markov Blanket framework for multimodal prognostic feature selection in TCGA-BRCA [8]. The present study does not introduce another feature-selection algorithm or a new prognostic signature. Instead, it asks which parts of a previously developed molecular modelling pipeline remain reproducible when moved from TCGA-BRCA to METABRIC.

We address this question using two complementary analysis tracks. Track A tests literal transport of historical TCGA-derived specifications. Molecular features are mapped to METABRIC without using METABRIC outcomes, and the source-trained feature sets, preprocessing specifications, and model coefficients are evaluated without target-cohort refitting. Track B asks a different question: whether the dependency-aware selection procedure can recover stable and prognostically useful molecular structure when reconstructed within METABRIC. All outcome-informed preprocessing, screening, feature selection, and model fitting are therefore restricted to outer-training folds under repeated outer cross-validation, with held-out folds used only for evaluation [9, 10].

In both tracks, molecular performance is interpreted relative to a matched clinical model. The primary quantity is the paired change in Harrell’s concordance index, Δ*C* = *C*_clinical+omics_ *− C*_clinical_, with both models evaluated on the same patients and under the same transport or resampling structure. Assayability, selection stability, gene or pathway recurrence, and incremental prognostic discrimination are reported separately; evidence in one dimension is not used as a substitute for performance in another.

After completion of the primary dual-track analyses, we added a protocol-locked post-hoc transport-decomposition analysis. The exact transportable historical RNA and copy-number panels were retained while model parameters were re-estimated locally within METABRIC. Size-matched random-panel benchmarks were then used to determine whether performance after local redevelopment was specific to the historical feature composition or was common among alternative assayable feature sets. Random panels were evaluated both against the broad METABRIC-assayable feature space and against the recovered historical candidate spaces from which the original modality-specific panels had been selected. These benchmarks were descriptive and were not treated as empirical-null hypothesis tests.

The resulting evaluation design separates five related but distinct questions: whether historical features are assayable in another cohort, whether a fixed source model transports, whether a fixed feature set retains information after local redevelopment, whether the selected features are enriched relative to reasonable alternatives, and whether independent reconstruction of the selection procedure yields stable and incrementally useful predictors. Overall survival is the primary endpoint, with recurrence-free survival evaluated as a prespecified Track B sensitivity endpoint.

## 2. Related Work

Breast-cancer prognostic modelling has long shown that molecular measurements can contribute to risk stratification, particularly through gene-expression signatures [1]. At the same time, high-dimensional molecular data contain extensive correlation and redundancy. Different feature sets may capture similar tumour structure, and random gene-expression signatures can also show apparent prognostic association [2]. This makes external evaluation important not only for confirming performance, but also for determining whether the exact selected representation carries information that persists outside the development cohort.

Large-scale benchmarking studies have reinforced this concern. Across multiple TCGA cancer datasets, multi-omics models have not consistently improved survival prediction over clinical models, despite access to substantially richer molecular information [6]. Standardized evaluations have likewise shown that reported performance depends strongly on the clinical baseline, modality availability, cohort composition, missing-data handling, and validation design [7]. These studies motivate direct comparison with a matched clinical model rather than interpretation of an omics-model C-index in isolation.

Another methodological issue is information leakage during molecular feature selection. In high-dimensional settings, screening or selecting features before cross-validation allows held-out outcome information to influence the fitted representation and can substantially bias performance estimates [9, 10]. Leakage-controlled resampling is therefore required when the feature-selection procedure itself is reconstructed in a new cohort, with all outcome-informed steps confined to the corresponding training data. This setting is fundamentally different from evaluation of a fixed historical model, where the source representation should be applied without outcome-guided modification.

Our previous work introduced a dependency-aware Markov Blanket framework for multimodal prognostic feature selection in TCGA-BRCA [8]. The present study focuses on a different problem: how to evaluate what remains reproducible when such a modelling pipeline is transferred to an independent cohort. Existing studies commonly report external model performance, feature recurrence, or within-cohort selection stability as separate analyses. Here, these quantities are placed within a single evaluation structure that distinguishes fixed-model transport, local redevelopment of the same feature set, independent reconstruction of the selection procedure, and incremental performance relative to matched clinical prediction.

## 3. Methods

### 3.1. Study design and evaluation tracks

We evaluated cross-cohort prognostic reproducibility using TCGA-BRCA as the source cohort and METABRIC as the independent target cohort [3, 4, 5]. The study was prognostic rather than causal: the objective was to determine whether molecular information improved survival discrimination beyond clinical information, not to estimate treatment effects.

The analysis was organized into two primary tracks. Track A evaluated literal transport of historical TCGA-derived prognostic specifications. Feature mapping was performed without using METABRIC outcomes, and the resulting source-trained models were applied without target-cohort coefficient refitting. Track B reconstructed the dependency-aware feature-selection procedure independently within METABRIC using repeated leakage-controlled outer cross-validation.

These tracks address different questions. Track A asks whether a fixed historical representation and its fitted risk function transport across cohorts. Track B asks whether the underlying selection procedure can identify stable and incrementally useful molecular structure when redeveloped in the target cohort. Assayability, selection stability, biological recurrence, and prognostic performance were therefore evaluated separately.

After completion of the primary Track A and Track B analyses, we performed a protocol-locked post-hoc extension to decompose the Track A transport result. This extension retained the same historical molecular features but allowed model parameters to be estimated locally within METABRIC, followed by size-matched random-panel benchmarks. These analyses were used to distinguish failure of literal model transport from loss of information in the fixed feature set and to contextualize the specificity of the historical feature composition. The primary study design is summarized in Fig. 1.

**Figure 1:**
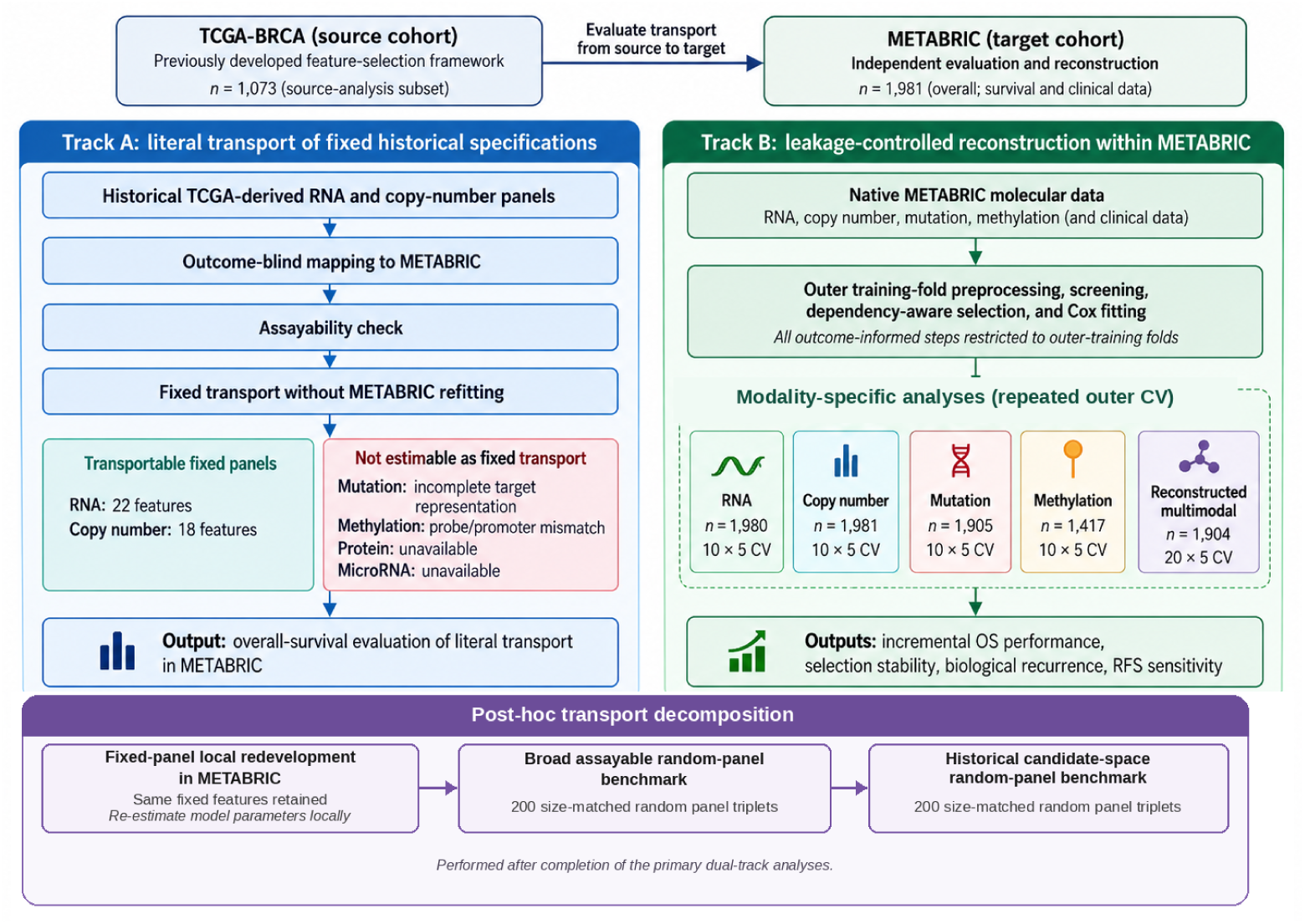
Primary dual-track study design. Track A evaluated outcome-blind transport of fixed TCGA-derived specifications to METABRIC without target-cohort refitting. Track B reconstructed the dependency-aware selection framework within METABRIC using leakage-controlled repeated cross-validation. The post-hoc fixed-panel redevelopment and random-panel benchmarks were performed after completion of these primary analyses.

### 3.2. Cohorts and endpoints

TCGA-BRCA provided the source molecular and clinical data used for the historical models. METABRIC provided the independent evaluation data. Molecular data included gene expression, copy-number alteration, somatic mutation, and methylation where available.

Overall survival (OS) was the primary endpoint. Survival time was measured in months from diagnosis to death or last recorded follow-up, with patients alive at last follow-up treated as censored. Analysis populations were defined according to endpoint availability and the molecular data required for each model.

Recurrence-free survival (RFS) was specified as a Track B sensitivity end-point. The Track B modelling procedure, outer-fold structure, repeat counts, candidate spaces, selector settings, Cox-model family, and seed sequences were retained from the corresponding OS analyses. All RFS-informed preprocessing, screening, selection, and fitting remained restricted to outer-training data. Because RFS was a sensitivity endpoint and several modality-specific contrasts were examined, its bootstrap intervals are reported without multiplicity adjustment and are interpreted as supporting rather than confirmatory evidence.

The primary OS analysis populations and resampling structures are summarized in Table 1. Endpoint-specific RFS populations are reported in the Supplementary Information.

**Table 1:** Overall-survival analysis populations and evaluation designs.

| Track | Analysis | Cohort | Evaluation design | $n$ |
| --- | --- | --- | --- | --- |
| A | Source specification | TCGA-BRCA | Fixed historical specification | 1,073 |
| A | Literal fixed-panel transport | METABRIC | No target-cohort refitting | 1,981 |
| B | RNA | METABRIC | 10 repeats × 5 folds | 1,980 |
| B | Copy number | METABRIC | 10 repeats × 5 folds | 1,981 |
| B | Mutation | METABRIC | 10 repeats × 5 folds | 1,905 |
| B | Methylation | METABRIC | 10 repeats × 5 folds | 1,417 |
| B | Reconstructed multimodal | METABRIC | 20 repeats × 5 folds | 1,904 |
| Post-hoc | Fixed-panel local redevelopment | METABRIC | 10 repeats × 5 folds | 1,980 |

### 3.3. Clinical comparators and molecular representation

Claims of molecular incremental value were made only relative to clinical models evaluated on the same patients and under the same analysis structure. Track A used the historical five-variable clinical specification comprising age, indicators for stage II, stage III, and stage IV disease, and an indicator of node-positive disease. Clinical imputation and scaling parameters were estimated in TCGA and transferred to METABRIC together with the source-model coefficients.

Track B used a harmonized METABRIC clinical panel consisting of age at diagnosis, grade, stage, tumour size, positive-node count, oestrogen-receptor status, progesterone-receptor status, HER2 status, and the Nottingham Prognostic Index [11]. Clinical preprocessing and model fitting were performed within each outer-training fold. Each modality-specific molecular model was compared with its fold-matched clinical model on the same modality-available population.

The secondary NPI analysis treated NPI alone as an additional clinical benchmark. It was performed after completion of the primary Track B analyses using locked out-of-fold predictions and common subsets with observed NPI values. NPI did not replace the primary multivariable clinical comparator, and no NPI-plus-omics model was fitted.

### 3.4. Track A: outcome-blind fixed-model transport

Track A evaluated the historical modality-specific feature selections from our previous TCGA-BRCA study [8]. A provenance audit linked the transported panels to the original modality-specific selected lists. The historical RNA selection contained 50 features from a 1,000-gene RNA candidate set (project identifier rna_8_composite_1000genes) and was obtained using the Incremental Association Markov Blanket (IAMB) algorithm with *α* = 0.05. The historical copy-number selection contained 50 features from a 1,000-gene copy-number candidate set (project identifier cnv_4_fdr_significant_ 1000genes) and was obtained using the Grow-Shrink Markov Blanket (GSMB) algorithm with *α* = 0.1.

TCGA and METABRIC used different identifiers and assay representations. Selected Ensembl identifiers were therefore mapped outcome-blindly using current and GRCh37 Ensembl annotations. Ambiguous or unresolved mappings were not resolved using METABRIC outcomes. A feature was retained only if the mapped gene was represented on the corresponding METABRIC platform.

This procedure retained 22 RNA and 18 copy-number features. Four fixed specifications were evaluated:

1. clinical variables only;
2. clinical variables plus the 22 transported RNA features;
3. clinical variables plus the 18 transported copy-number features; and
4. clinical variables plus both transported molecular panels.

For RNA, the source and target measurements were transformed separately using an outcome-blind rank-based normal-score transformation. For copy number and clinical variables, imputation and scaling parameters were estimated from the TCGA source data and applied to METABRIC. Cox-model coefficients were estimated in TCGA and transferred unchanged. METABRIC survival outcomes were therefore used only to evaluate the transported risk scores.

Track A was treated as external evaluation of a fixed Cox prognostic specification rather than as target-cohort model redevelopment [12].

Exact fixed-panel transport was not attempted for modalities lacking a comparable target representation. The available METABRIC mutation data did not support complete one-to-one representation of the historical mutation panel. Similarly, the historical TCGA methylation features and the METABRIC promoter-level measurements did not provide a sufficiently comparable feature space for literal transport. Compatible protein and microRNA layers were unavailable. These limitations apply to fixed-model transport only and did not prevent independent within-METABRIC analysis in Track B.

### 3.5. Track B: leakage-controlled reconstruction

Track B evaluated whether the dependency-aware feature-selection framework could recover useful structure independently within METABRIC. It did not transport the historical TCGA feature list.

The historical conditional-independence software could not be reproduced bitwise in the current environment. We therefore used a locked reconstructed implementation based on rank-Gaussian transformations and partial-association testing while retaining the forward-inclusion and backward-removal structure of the IAMB family of Markov Blanket procedures [13, 8]. The analysis is therefore described as a reconstructed methodological replication rather than an exact software reproduction.

RNA, copy number, mutation, and methylation were evaluated separately using 10 repetitions of five-fold outer cross-validation. The reconstructed multimodal analysis used 20 repetitions of five-fold outer cross-validation. All outcome-informed steps were confined to the corresponding outer-training fold, including molecular preprocessing, supervised candidate screening, dependency-aware selection, clinical preprocessing, and Cox-model fitting. Held-out observations were used only for prediction. This design prevents outcome information from entering feature selection before performance evaluation [9, 10].

For continuous molecular modalities, candidate screening was performed within the training fold using signed Spearman association with survival time, retaining up to the 100 strongest positive and 100 strongest negative associations before dependency-aware selection. Mutation candidates were restricted to nonsynonymous genes represented in the 173-gene METABRIC mutation panel and required a training-fold mutation frequency of at least 1%; candidate sets were capped at 200 features. All selector thresholds and modality-specific settings were fixed before the full repeated analyses.

For each modality-specific outer fit, three penalized Cox models were evaluated: clinical only, selected molecular features only, and clinical plus selected molecular features. Cox fitting used a fixed convergence sequence of penalization values 0.05, 0.2, and 1.0; the first value yielding a successful fit was used. This sequence was a convergence rule rather than performance-based tuning.

Each patient received one out-of-fold prediction per repeat from a model for which that patient had not contributed to preprocessing, feature screening, selection, or fitting. Selected feature sets were retained for each outer fit. Selection reproducibility was summarized using selection frequency, pairwise Jaccard similarity, overlap coefficients, and chance-adjusted overlap, which corrects pairwise overlap for the overlap expected from the candidate-set size. Exact gene overlap between the assayable historical TCGA panels and recurrent METABRIC selections was used to summarize gene-level recurrence. Pathway-level recurrence was treated descriptively by comparing top-ranked pathway sets between cohort-specific analyses and by checking whether any pathway met the false-discovery-rate criterion used in the original analysis [8] in both cohorts. These recurrence summaries were evaluated separately from predictive performance.

### 3.6. Post-hoc fixed-panel redevelopment and random-panel benchmarks

The post-hoc extension was designed and locked after completion of the primary dual-track analyses. It did not modify the historical Track A results.

First, we evaluated whether the fixed Track A feature sets retained prognostic information when model parameters were allowed to adapt to METABRIC. The 22 RNA and 18 copy-number features were kept unchanged; no molecular feature selection was performed. A common OS cohort containing patients represented in both fixed molecular matrices was used for all four model specifications: clinical only, clinical plus RNA, clinical plus copy number, and clinical plus RNA and copy number.

Local redevelopment used 10 repetitions of five-fold outer cross-validation, with identical folds for all four specifications. Clinical and copy-number imputation and scaling were estimated using outer-training data only. RNA was transformed using an empirical-CDF normal-score transformation estimated from the outer-training fold; held-out RNA values were mapped only through the training-fold empirical distributions.

All four models within a fold were required to use the same Cox penalization value. Penalization values of 0.05, 0.2, and 1.0 were considered in that order, advancing to the next value only if at least one of the four models failed to fit. This produced a paired comparison in which differences between clinical-only and molecular models could not be attributed to different regularization choices.

We then performed two descriptive size-matched random-panel benchmarks. Each benchmark used 200 pre-generated random panel pairs containing 22 RNA and 18 copy-number features. Random-panel identities were generated and hashed before model fitting, and the exact METABRIC cohort, outer folds, clinical baseline, preprocessing rules, and Cox penalization used in the fixed-panel redevelopment were retained.

The first benchmark sampled features from a broad METABRIC-assayable universe. Eligible features were required to occur in the corresponding METABRIC molecular matrix, have at least 95% non-missing measurements, and have non-zero variance greater than 10*^−^*^12^. Historical features were not removed from the sampling universe.

The second benchmark restricted random sampling to the recovered historical candidate spaces from which the original modality-specific selections had been made. The corresponding historical RNA and copy-number candidate matrices each contained 1,000 features. Candidate identifiers were mapped to METABRIC outcome-blindly using locally available mappings and, where necessary, current and GRCh37 Ensembl annotations. Candidates were then intersected with the previously locked METABRIC-assayable universe. This produced assayable historical candidate spaces of 555 RNA genes and 350 copy-number genes.

For both benchmarks, performance was summarized using fold-local Harrell C-index contrasts. Within each repeat, fold-level Δ*C* values were weighted by held-out fold size and then averaged across the five folds; the resulting repeat-level contrasts were averaged across the 10 repeats. Historical panel performance was compared with the distribution of the 200 size-matched random panels using descriptive medians, quantiles, ranks, and percentiles. These random-panel distributions were not treated as empirical null distributions, and their ranks or exceedance counts were not interpreted as *p*-values.

### 3.7. Performance measures and uncertainty

The primary discrimination measure was Harrell’s concordance index [14]. Incremental molecular performance was defined as

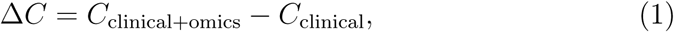

where both models were evaluated on the same patients under the corresponding transport or resampling design.

Five-year discrimination was evaluated as a secondary measure using censoring-aware cumulative/dynamic AUC [15]. Censoring was handled using inverse-probability-of-censoring weights (IPCW). Track A additionally retained Uno’s C-index and prediction-error measures, including Brier scores and the integrated Brier score, as supporting diagnostics [16, 17]. These secondary metrics were not used to override conclusions based on the primary Harrell C-index contrast.

For Track A, uncertainty in the fixed transported predictions was estimated using 1,000 paired patient-bootstrap samples. Clinical-only and molecular risks were evaluated on the same resampled patients in each bootstrap sample. For Track B, 2,000 paired patient-bootstrap samples were drawn from the locked repeated out-of-fold predictions. The same sampled patients were used for the clinical and corresponding molecular model within each bootstrap replicate, and performance contrasts were averaged across repeats. The same procedure was used for the RFS sensitivity analysis and the secondary NPI comparison.

The fixed-panel local-redevelopment analysis likewise used 2,000 paired patient bootstrap samples drawn only after the repeated out-of-fold predictions had been completed and hash-locked. These bootstrap intervals quantify uncertainty conditional on the fitted repeated models. They are not full-pipeline bootstrap intervals because preprocessing, model fitting, and, for Track B, feature selection were not repeated inside each bootstrap sample.

For the primary OS contrasts, a molecular model was described as showing positive incremental performance when the entire 95% paired-bootstrap interval for Δ*C* was above zero and negative incremental performance when the entire interval was below zero. Intervals crossing zero were described as showing no reliable incremental improvement. This classification rule does not constitute an equivalence test; an interval crossing zero was not interpreted as evidence that two models were equivalent.

Repeated-split distributions and feature-stability summaries were interpreted as measures of algorithmic variability rather than population confidence intervals. The two random-panel benchmark analyses were descriptive contextualization analyses only and did not contribute additional hypothesis tests. Model performance was assessed across complementary dimensions of discrimination and prediction error [18].

### 3.8. Software

All analyses were executed locally using version-controlled Python scripts. Analysis code and configuration files are available in the public repository described in the Code availability statement.

## 4. Results

### 4.1. Assayability and analysis populations

The fixed Track A specifications were evaluated in 1,981 METABRIC patients with available overall-survival and clinical data. Outcome-blind harmonization retained 22 of the historical RNA features and 18 of the historical copy-number features.

Exact transport was more limited for the remaining modalities. The available METABRIC mutation data did not provide complete representation of the historical mutation panel. For methylation, the historical TCGA probe-level features could not be represented reliably in the METABRIC promoter-level feature space. These modalities were therefore not classified as failed biological replications; rather, exact fixed-panel transport was not estimable.

Track B used METABRIC’s native molecular representations. The primary modality-specific OS analyses included 1,980 patients for RNA, 1,981 for copy number, 1,905 for mutation, and 1,417 for methylation. The reconstructed multimodal analysis included 1,904 patients. Analysis populations and resampling structures are summarized in Table 1.

### 4.2. Track A: literal transport of the fixed TCGA specifications

The transported five-variable clinical model achieved a Harrell C-index of 0.6212 in METABRIC. Adding the fixed copy-number panel produced a C-index of 0.6215, whereas the clinical-plus-RNA and clinical-plus-RNA-plus-copy-number models achieved C-indices of 0.6076 and 0.6091, respectively.

Using the unrounded point estimates, the copy-number panel changed discrimination by only Δ*C* = 0.0003 (95% paired-bootstrap interval, *−*0.0034 to 0.0042). The transported RNA panel reduced discrimination by Δ*C* = *−*0.0136 (95% interval, *−*0.0262 to *−*0.0002). The combined RNA and copy-number model also had a negative point contrast, Δ*C* = *−*0.0121, although its interval crossed zero (95% interval, *−*0.0254 to 0.0018).

Supporting measures did not indicate a clear compensating advantage. Five-year censoring-aware AUC values were 0.6115 for clinical only, 0.6110 for clinical plus RNA, 0.6136 for clinical plus copy number, and 0.6161 for the combined model. Integrated Brier scores varied little, from 0.1547 to 0.1554. Calibration slopes were 1.247, 0.814, 1.242, and 0.803 for the clinical, RNA, copy-number, and combined specifications, respectively. Thus, the RNA-containing models also departed further from the ideal calibration slope of one.

These results show that successful cross-platform representation of the historical features was not sufficient for successful transport of the source-trained prognostic models.

### 4.3. Post-hoc transport decomposition

We next asked whether the same fixed molecular features retained prognostic information when their coefficients were re-estimated within METABRIC. This post-hoc analysis used a common cohort of 1,980 patients, including 1,143 deaths, and 10 repetitions of five-fold outer cross-validation. The molecular feature sets remained fixed at 22 RNA and 18 copy-number features; no feature selection was repeated.

Local redevelopment changed the Track A result substantially. The mean repeated out-of-fold clinical C-index was 0.6237. Clinical plus fixed RNA achieved a C-index of 0.6400 (Δ*C* = 0.0162; 95% conditional bootstrap interval, 0.0076–0.0244), while clinical plus fixed copy number achieved 0.6328 (Δ*C* = 0.0091; 95% interval, 0.0031–0.0149). The combined fixed RNA and copy-number model achieved 0.6438 (Δ*C* = 0.0201; 95% interval, 0.0103–0.0298).

The corresponding fold-contained five-year IPCW AUC gains were 0.0307 for RNA, 0.0185 for copy number, and 0.0392 for the combined panel. All 10 repeat-level Δ*C* estimates were positive for each of the three molecular specifications. The prespecified penalizer of 0.05 was sufficient for all 50 outer folds, so no convergence fallback was required.

The contrast with literal transport was therefore substantial. The same RNA features changed from a direct Track A Δ*C* of *−*0.0136 to +0.0162 after local redevelopment, while the combined panel changed from *−*0.0121 to +0.0201. These differences are descriptive rather than paired inferential contrasts because the literal-transport and local-redevelopment analyses used different evaluation designs.

#### 4.3.1. Size-matched random-panel contextualization

The positive local-redevelopment results were then compared with size-matched random panels using the same cohort, folds, preprocessing, clinical baseline, and Cox-model specification.

In the broad assayable benchmark, the eligible feature spaces contained 20,385 RNA and 22,542 copy-number genes. The historical fixed panels were above the median random-panel performance but were not extreme. On the fold-local scale used for this comparison, the historical RNA panel had Δ*C* = 0.0162 and was at the 68.5th percentile of 200 random 22-gene RNA panels; 63 of 200 random panels matched or exceeded its performance. The historical copy-number panel was at the 64.5th percentile, with 71 of 200 random panels matching or exceeding it. The combined panel was more highly ranked, at the 83.0th percentile, although 34 of 200 random panel pairs still matched or exceeded the historical result.

Positive incremental discrimination was common in this broad benchmark. All 200 random RNA panels, 190 of 200 copy-number panels, and 199 of 200 combined panels produced positive mean Δ*C* values after local redevelopment.

We then restricted sampling to the recovered historical candidate spaces. Each original modality-specific candidate matrix contained 1,000 features. After outcome-blind mapping and intersection with the METABRIC-assayable universe, 555 RNA and 350 copy-number candidate genes remained.

The historical RNA panel remained only moderately ranked within its own candidate space: its percentile was 65.0, and 70 of 200 random RNA panels matched or exceeded its performance. In contrast, the historical copy-number panel exceeded all 200 size-matched random panels drawn from its recovered assayable candidate space. The historical combined panel was at the 81.5th percentile, with 37 of 200 random panel pairs matching or exceeding it.

The random-panel results are summarized together with the fixed-panel redevelopment estimates in Table 2 and visualized in Fig. 4. The random-panel ranks and percentiles are descriptive benchmarks and are not interpreted as *p*-values.

**Table 2:** Post-hoc local redevelopment of the fixed historical panels and size-matched random-panel contextualization. Local-redevelopment intervals are conditional paired patient-bootstrap intervals from locked repeated out-of-fold predictions. Random-panel percentiles are descriptive and are not *p*-values.

| Fixed panel | Local $\Delta C$ | 95% interval | IPCW $\Delta AUC_{5y}$ | Broad percentile | Candidate-space percentile |
| --- | --- | --- | --- | --- | --- |
| RNA | 0.0162 | [0.0076, 0.0244] | 0.0307 | 68.5 | 65.0 |
| Copy number | 0.0091 | [0.0031, 0.0149] | 0.0185 | 64.5 | 100.0 |
| RNA + copy number | 0.0201 | [0.0103, 0.0298] | 0.0392 | 83.0 | 81.5 |

### 4.4. Track B: leakage-controlled prognostic performance

Independent feature selection within METABRIC did not produce reliable OS improvement for RNA, copy number, or mutation. Relative to their fold-matched clinical models, the paired contrasts were Δ*C* = 0.0001 for RNA (95% interval, *−*0.0065 to 0.0071), Δ*C* = 0.0024 for copy number (95% interval, *−*0.0025 to 0.0077), and Δ*C* = *−*0.0014 for mutation (95% interval, *−*0.0072 to 0.0041).

Methylation performed worse than its clinical comparator (Δ*C* = *−*0.0217; 95% interval, *−*0.0314 to *−*0.0125). The reconstructed multimodal model also showed a negative contrast (Δ*C* = *−*0.0128; 95% interval, *−*0.0233 to *−*0.0023).

The absolute and incremental OS estimates are summarized in Table 3 and Fig. 2. Track A and Track B therefore produced different forms of negative evidence: literal transport failed for the RNA-containing historical models, whereas independent target-cohort feature selection did not recover reliable OS improvement over the matched clinical models.

**Figure 2:**
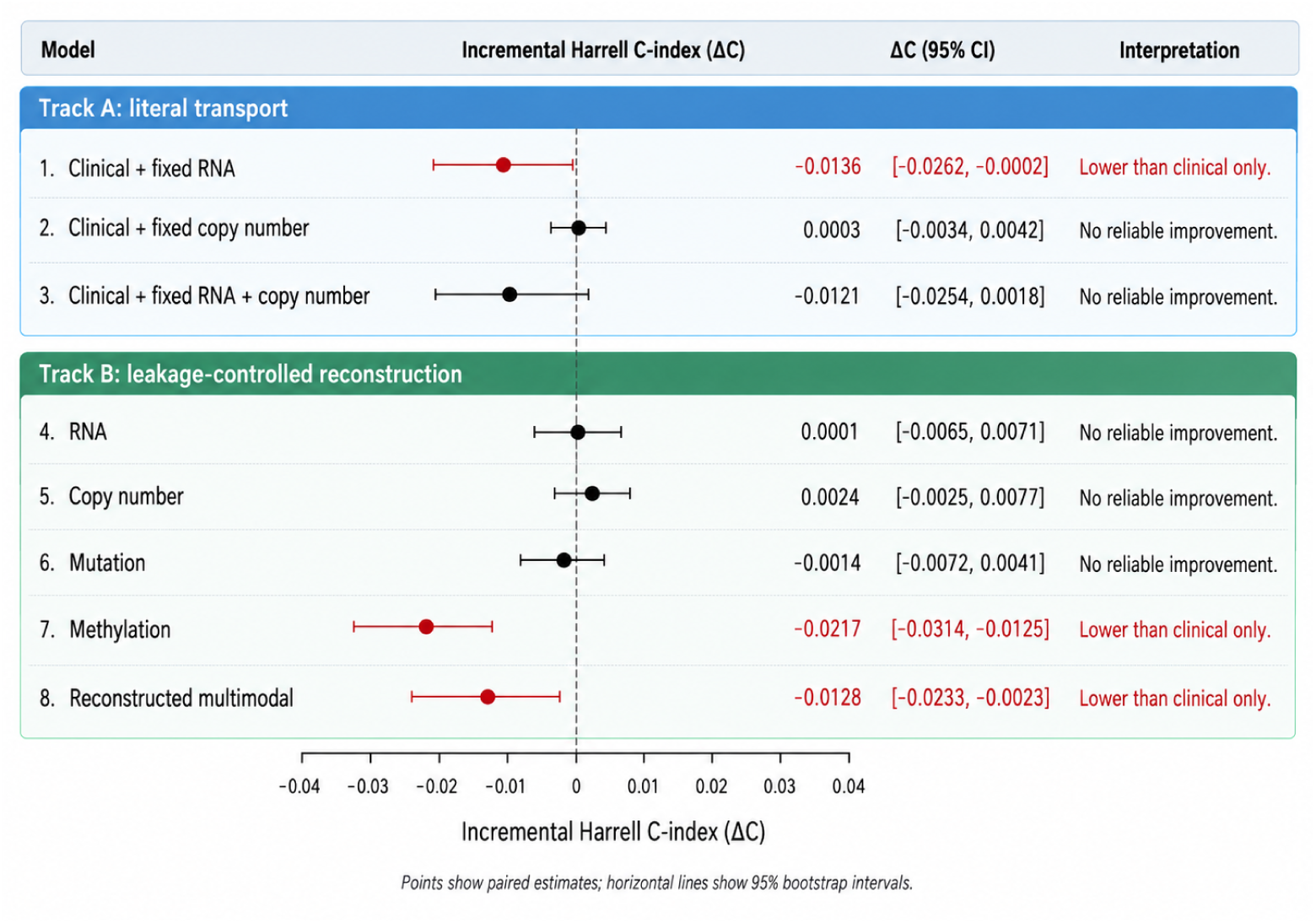
Incremental overall-survival discrimination relative to matched clinical models. Points show the paired change in Harrell’s C-index and horizontal lines show 95% paired patient-bootstrap intervals. Track A evaluates literal fixed-model transport, whereas Track B evaluates leakage-controlled reconstruction within METABRIC.

**Figure 3:**
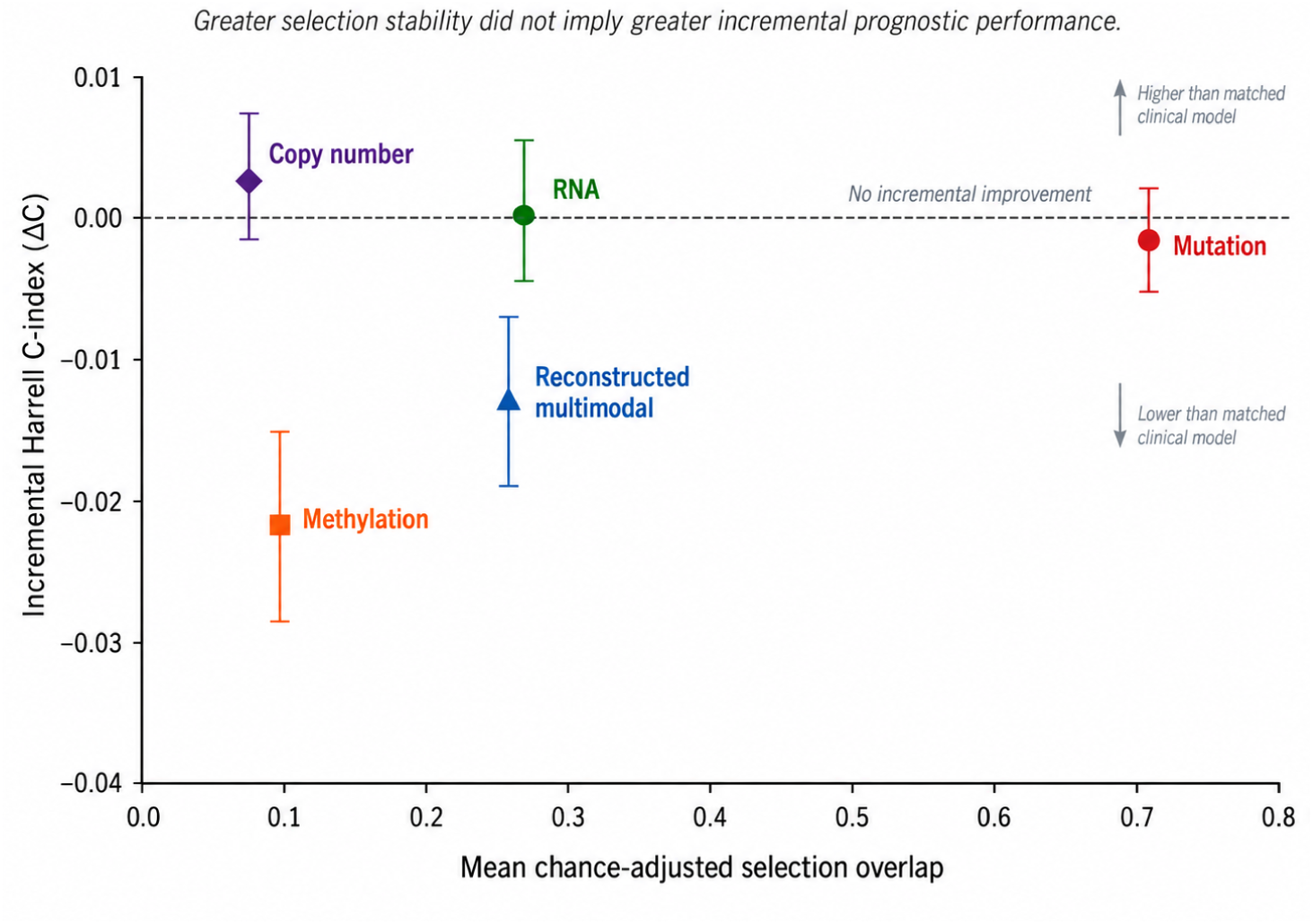
Selection stability and incremental overall-survival performance in Track B. The horizontal axis shows mean chance-adjusted overlap across outer-training fits. The vertical axis shows the paired incremental Harrell C-index relative to the fold-matched clinical model. Greater selection stability did not imply greater incremental prognostic performance.

**Figure 4:**
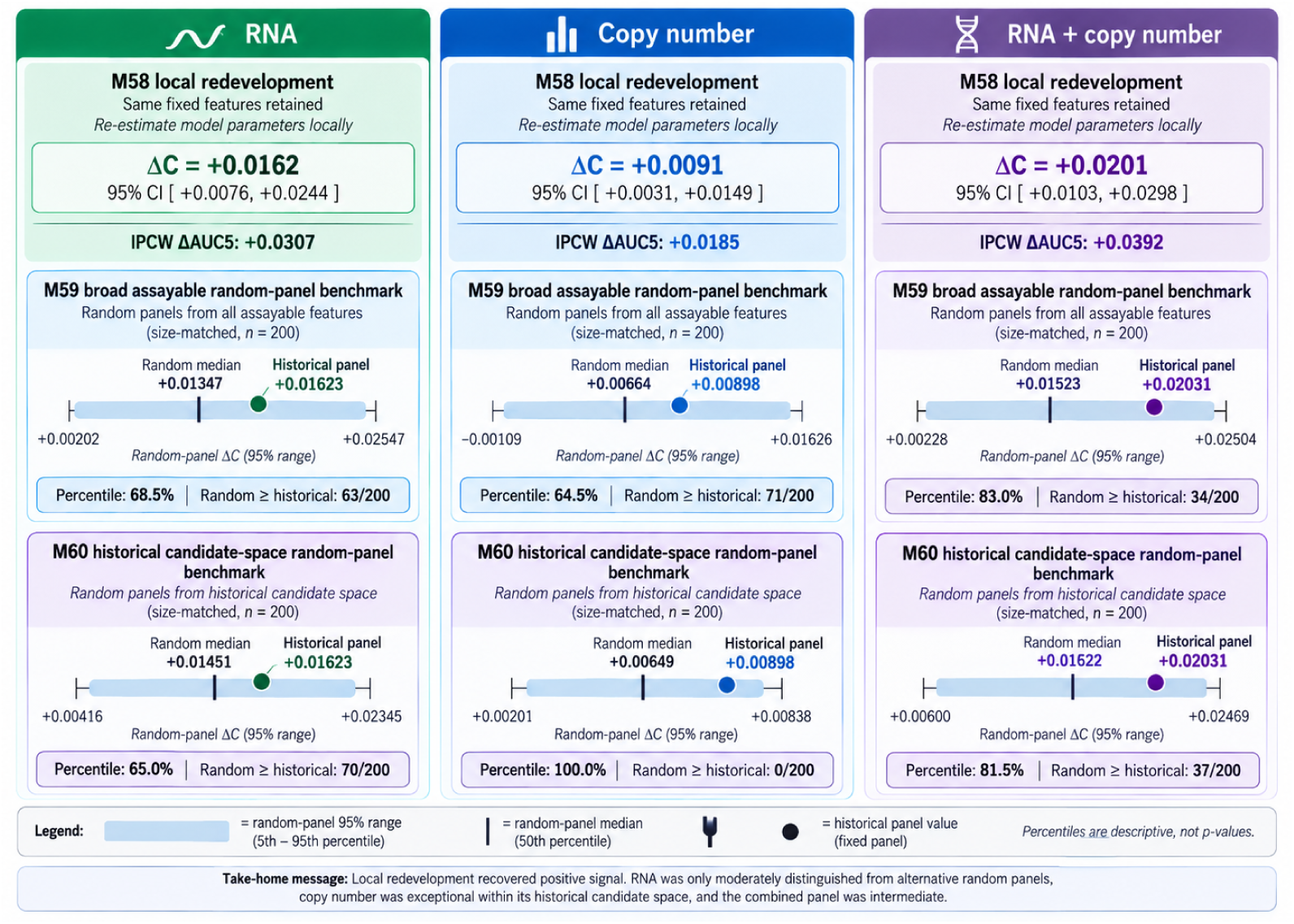
Fixed-panel local redevelopment and size-matched random-panel contextualization. For each molecular specification, the upper panel shows the incremental overall-survival discrimination obtained after local METABRIC redevelopment of the fixed historical feature set. The lower panels compare the historical fixed-panel performance with 200 size-matched random panels sampled from (i) the broad METABRIC-assayable feature universe and (ii) the recovered assayable historical candidate space. Random-panel medians and descriptive 95% ranges are shown together with the historical panel value and its descriptive percentile. Labels M58–M60 identify the three locked post-hoc analysis stages shown in the figure. The benchmark statistic uses the fold-local weighted contrast defined in Methods, so historical values used for percentile placement can differ slightly from the patient-level redevelopment estimates shown in the upper panels. Random-panel distributions were used for contextualization only and were not interpreted as empirical null distributions or *p*-values.

**Table 3:** Overall-survival discrimination and incremental molecular performance. Track A values are fixed external-evaluation estimates; Track B values are mean repeated out-of-fold estimates. Paired intervals were calculated from unrounded values.

| Molecular specification | Clinical C | Clinical + omics C | $\Delta C$ (95% interval) | Interpretation |
| --- | --- | --- | --- | --- |
| *Track A: literal fixed-model transport* |  |  |  |  |
| Copy number | 0.6212 | 0.6215 | 0.0003 [−0.0034, 0.0042] | No reliable improvement |
| RNA | 0.6212 | 0.6076 | −0.0136 [−0.0262, −0.0002] | Lower than clinical only |
| RNA + copy number | 0.6212 | 0.6091 | −0.0121 [−0.0254, 0.0018] | No reliable improvement |
| *Track B: leakage-controlled reconstruction* |  |  |  |  |
| RNA | 0.6693 | 0.6694 | 0.0001 [−0.0065, 0.0071] | No reliable improvement |
| Copy number | 0.6689 | 0.6713 | 0.0024 [−0.0025, 0.0077] | No reliable improvement |
| Mutation | 0.6690 | 0.6677 | −0.0014 [−0.0072, 0.0041] | No reliable improvement |
| Methylation | 0.6735 | 0.6517 | −0.0217 [−0.0314, −0.0125] | Lower than clinical only |
| Reconstructed multimodal | 0.6692 | 0.6564 | −0.0128 [−0.0233, −0.0023] | Lower than clinical only |

In the secondary NPI comparison, the multivariable clinical-only models outperformed NPI by approximately 0.035–0.037 C-index units across the analysis-specific common subsets. In the 1,904-patient multimodal subset, clinical plus omics exceeded NPI by 0.0237 (95% interval, 0.0060–0.0420) but remained below the matched clinical-only model by 0.0129 (95% interval, *−*0.0239 to *−*0.0022). Thus, exceeding NPI did not imply incremental molecular performance over the primary clinical comparator.

### 4.5. Recurrence-free-survival sensitivity analysis

The RFS sensitivity analysis used endpoint- and modality-specific populations: 1,979 patients with 803 recurrence events for RNA, 2,078 with 832 events for copy number, 2,304 with 932 events for mutation, 1,418 with 580 events for methylation, and 1,903 with 771 events for the reconstructed multimodal analysis.

RNA showed positive incremental RFS discrimination. The clinical-only and clinical-plus-RNA C-indices were 0.6496 and 0.6642, respectively, giving Δ*C* = 0.0146 with a 95% conditional bootstrap interval of 0.0035–0.0258.

Copy number showed a smaller positive point estimate (Δ*C* = 0.0059; 95% interval, *−*0.0026 to 0.0144), while mutation showed Δ*C* = *−*0.0040 (95% interval, *−*0.0107 to 0.0030). Methylation remained below its clinical comparator (Δ*C* = *−*0.0170; 95% interval, *−*0.0319 to *−*0.0017). The reconstructed multimodal model showed Δ*C* = 0.0029 (95% interval, *−*0.0136 to 0.0204).

The RNA result was therefore specific to the RFS sensitivity analysis and did not extend to the reconstructed multimodal model. Because RFS was a secondary endpoint and several modality-specific contrasts were evaluated, these unadjusted intervals are interpreted as sensitivity evidence rather than as independent confirmatory tests. Complete RFS results are reported in Supplementary Table S1.

### 4.6. Selection stability and biological recurrence

Feature-selection stability differed markedly across modalities. Mutation showed the highest mean chance-adjusted overlap across outer-training fits (0.715). RNA and the reconstructed multimodal analysis showed intermediate values of 0.288 and 0.285, respectively, whereas stability was lower for methylation (0.109) and copy number (0.089).

These stability estimates did not track incremental OS discrimination. Mutation was the most reproducibly selected modality but had Δ*C* = *−*0.0014. RNA showed moderate selection stability with essentially zero Track B OS improvement, and the reconstructed multimodal analysis combined moderate stability with a negative performance contrast.

Exact gene-level recurrence between the assayable historical TCGA panels and the recurrent METABRIC selections was not observed for RNA, copy number, or mutation. Gene-level methylation comparison was not estimable because the historical probe representation could not be mapped adequately to the METABRIC promoter-level feature space.

Pathway-level recurrence was limited and descriptive. Mutation shared six of its top 20 pathways between the cohort-specific analyses, while the pooled comparison shared five. No pathway met the study’s false-discovery-rate criterion in both cohorts. Stability and biological recurrence therefore provided information about reproducibility of the selected representation but did not alter the paired prognostic-performance conclusions.

## 5. Discussion

### 5.1. Principal findings

This study shows that cross-cohort reproducibility of a molecular prognostic model cannot be described adequately by a single external-validation result. Different components of the modelling pipeline behaved differently when moved from TCGA-BRCA to METABRIC.

Literal transport in Track A was weak. The fixed copy-number specification was essentially neutral relative to the transported clinical model, while the RNA-containing models showed lower discrimination. However, this result did not imply that the transported molecular features themselves lacked prognostic information. When the exact same 22 RNA and 18 copy-number features were retained but model parameters were re-estimated locally within METABRIC, incremental discrimination became positive for RNA, copy number, and their combination. The largest gain was observed for the combined fixed panel (Δ*C* = 0.0201).

Calibration and discrimination capture different aspects of model performance and may deteriorate differently under dataset shift [19].

The random-panel benchmarks provided an important qualification. In the broad METABRIC-assayable feature space, locally refitted random panels frequently improved discrimination as well. The historical RNA and copy-number panels ranked only at the 68.5th and 64.5th percentiles, respectively, and the combined panel at the 83.0th percentile. Thus, the positive local-redevelopment result could not be attributed simply to unique information carried by the exact historical feature composition.

Restricting the benchmark to the recovered historical candidate spaces revealed a modality-specific pattern. The RNA panel remained only moderately ranked (65th percentile), whereas the historical copy-number panel exceeded all 200 size-matched random copy-number panels sampled from its assayable candidate space. The combined panel remained moderately high but not extreme (81.5th percentile). These results suggest that local model adaptation and feature-set specificity contributed differently across molecular modalities.

Track B provided a complementary result. Independent feature selection within METABRIC did not produce reliable OS improvement for RNA, copy number, or mutation, while methylation and the reconstructed multimodal model performed worse than their matched clinical comparators. Selection stability also failed to predict incremental performance: mutation was the most stable modality but provided essentially no additional discrimination.

Taken together, these findings separate at least three distinct forms of reproducibility: transport of a fitted source model, retention of prognostic information in a fixed feature set after local redevelopment, and reproducibility of a feature-selection procedure. Success in one did not imply success in the others.

### 5.2. Model transport and local redevelopment

The contrast between literal transport and local redevelopment is one of the most informative findings of the study. A fixed model may fail in a new cohort because the relationship between its predictors and outcome differs across populations, because predictor distributions shift, because preprocessing does not transfer perfectly, or because the source coefficients are poorly calibrated to the target population [20, 21]. These mechanisms cannot be distinguished from a single external C-index alone.

Our results show that the negative Track A RNA result should not be interpreted as evidence that the underlying feature set was devoid of prognostic information. The same RNA features produced positive incremental discrimination once parameters were estimated within METABRIC. The combined panel showed an even larger change, from a negative point contrast under literal transport to positive incremental performance after local redevelopment.

At the same time, this comparison should not be reduced to a claim of “coefficient transport failure.” The local-redevelopment analysis changed the estimation setting as a whole: model parameters were learned in METABRIC, preprocessing was confined to training folds, and performance was evaluated using repeated out-of-fold prediction. The data therefore support the broader conclusion that the historical feature sets contained locally usable information despite poor transport of the complete source-trained specifications.

This distinction is relevant beyond the present cohorts. External validation, model updating, and redevelopment answer different questions and should be reported separately rather than treated as interchangeable evidence of generalizability [21]. For high-dimensional omics models, where cohort and platform shift may affect both predictor distributions and fitted effects, reporting only the performance of the unmodified source model can obscure whether failure arises from the representation itself or from the way that representation was parameterized.

### 5.3. How specific were the historical feature sets?

The size-matched benchmarks show why local redevelopment alone is not sufficient to establish that a historical molecular signature is uniquely informative. This issue is particularly relevant in gene-expression data, where correlated molecular programs allow many different feature sets to encode related tumour biology [2]. Large multi-omics survival benchmarks have also shown that predictive gains vary substantially with cohort, modality, and evaluation design [6, 7].

In the broad assayable benchmark, almost every random RNA panel and nearly every combined random panel showed a positive mean incremental C-index after local fitting. The historical RNA panel was therefore better than the median random panel but far from exceptional. The historical candidate-space benchmark reached the same conclusion within the RNA candidate space: 70 of 200 random candidate-derived RNA panels matched or exceeded the historical panel. For RNA, the present results therefore provide little evidence that the exact selected feature identities were the main reason local redevelopment succeeded.

Copy number behaved differently. In the broad assayable universe, the historical copy-number panel was only moderately ranked. When sampling was restricted to the recovered historical candidate space, however, none of the 200 random 18-feature panels exceeded the historical panel. This result is consistent with enrichment of useful copy-number features by the historical selection procedure within its own candidate space.

The CNA benchmark should nevertheless be interpreted cautiously. The numerical difference between the historical panel and the best sampled alternative was small, only 200 random panels were evaluated, and the benchmark distribution was specified as descriptive rather than as an empirical null distribution. The result therefore supports relative enrichment within the evaluated candidate space but does not establish statistical significance, uniqueness, or biological causality.

The combined panel occupied an intermediate position. It ranked above most random panels in both benchmarks but was not extreme. This is compatible with a model in which the combined signal reflects both a comparatively specific copy-number component and a broader RNA component for which many alternative feature subsets retain similar prognostic information.

### 5.4. Selection stability and biological recurrence

Track B illustrates a separate limitation of molecular reproducibility: features can be selected consistently without improving prediction beyond clinical information. Mutation had by far the highest chance-adjusted selection overlap, yet its incremental OS C-index was close to zero. RNA and the reconstructed multimodal analysis showed moderate stability without reliable OS improvement, while copy number had low stability despite a small positive point estimate.

Selection stability should therefore be viewed as a property of the modelling procedure rather than as a surrogate for prognostic value. Stable features may represent reproducible molecular structure, but their information can be redundant with clinical predictors or too weak to improve patient ranking. Conversely, unstable individual features can arise when several correlated variables capture similar underlying biology.

The limited cross-cohort gene recurrence is consistent with this interpretation. Exact gene overlap between historical TCGA panels and recurrent METABRIC selections was absent for RNA, copy number, and mutation. Pathway-level overlap was somewhat greater but remained descriptive, and no pathway met the prespecified false-discovery threshold in both cohorts. Biological recurrence, selection stability, and predictive utility therefore provided complementary rather than interchangeable evidence.

### 5.5. Endpoint dependence

The RFS sensitivity analysis differed from the primary OS results. RNA improved RFS discrimination by approximately 0.015 C-index units, whereas its Track B OS contrast was essentially zero. Methylation remained below the clinical comparator for both endpoints, while copy number, mutation, and the reconstructed multimodal model showed no reliable RFS improvement.

This endpoint-specific RNA result is potentially informative but should not be overinterpreted. RFS was a secondary sensitivity endpoint, several modality-specific contrasts were examined, and the reported bootstrap intervals were not adjusted for multiplicity. In addition, OS and RFS analyses did not use identical patient populations for every modality. The positive RNA RFS contrast therefore provides evidence of possible endpoint dependence rather than confirmation of a general RNA prognostic advantage.

Independent replication would be required to determine whether the RFS finding reflects a reproducible difference between recurrence and mortality prediction or sampling variation in the present cohort.

### 5.6. Methodological implications

The results support a staged approach to external evaluation of high-dimensional prognostic models.

First, assayability should be established before predictive transport is interpreted. Failure to represent a source feature on a target platform is different from failure of a measurable feature to reproduce.

Second, literal transport of a fitted model should be distinguished from local redevelopment of its fixed feature set. A negative external result does not by itself determine whether the representation or the fitted model is responsible.

Third, positive performance after local redevelopment should be contextualized against reasonable alternative feature sets. In high-dimensional molecular data, a locally fitted panel may improve prediction even when its exact composition is not special. Size-matched random-panel benchmarking provided a simple way to expose this distinction in the present study.

Finally, reconstruction of the original feature-selection procedure addresses a different question again. Track B tested whether the dependency-aware framework could identify stable and useful structure in METABRIC, not whether the historical TCGA feature list transported. Keeping these analyses separate made it possible to identify different failure modes that would otherwise have been collapsed into a single external-validation conclusion.

Across all stages, incremental molecular performance was evaluated against a clinical comparator on the same patients and within the same analysis structure. This paired design is important because an apparently reasonable absolute C-index for a molecular model does not establish that the molecular features add information beyond clinical prediction [22].

### 5.7. Strengths and limitations

A major strength of the study is the separation of assayability, literal model transport, fixed-feature local redevelopment, selection-framework reconstruction, feature stability, biological recurrence, and incremental performance. Track A mapping was outcome-blind, Track B confined outcome-informed operations to outer-training folds, and the post-hoc redevelopment and random-panel analyses were protocol-locked before model fitting. The same matched-comparator principle was retained throughout.

The provenance audit also established the relationship between the transported features and the historical modality-specific selections. This allowed the candidate-space benchmark to use the actual recovered RNA and copy-number candidate datasets rather than an arbitrarily chosen gene universe.

Several limitations remain. First, METABRIC was the only principal external cohort. The extent to which the observed transport and redevelopment patterns generalize to other populations, platforms, and clinical settings therefore remains unknown.

Second, only assayable subsets of the historical RNA and copy-number panels could be evaluated under literal transport. Exact transport of the mutation and methylation representations was not possible, and comparable protein and microRNA data were unavailable. Track A consequently evaluates the transportable portion of the historical molecular representation rather than the complete original multimodal model.

Third, the post-hoc fixed-panel redevelopment and random-panel benchmark analyses were motivated by the observed Track A results. Although their protocols, random panels, and analysis settings were locked before the corresponding model fits, they should be interpreted as mechanistic follow-up analyses rather than as prespecified confirmatory tests.

Fourth, the random-panel benchmarks used 200 panels per analysis. Their percentiles and exceedance counts provide useful descriptive context but are not formal tail-probability estimates. In particular, the observation that no candidate-space random CNA panel exceeded the historical CNA panel should not be converted retrospectively into a significance test.

Fifth, the bootstrap intervals used for repeated Track B and local redevelopment analyses were conditional on the fitted out-of-fold models. They did not repeat feature selection, preprocessing, or model estimation inside each bootstrap sample and therefore do not capture full model-development uncertainty.

Finally, the OS and RFS analyses used endpoint- and modality-specific populations, and the positive RNA RFS result was one of several secondary contrasts. These findings require independent replication before being interpreted as endpoint-specific molecular prognostic effects. The present study is prognostic and does not support causal, treatment-effect, or biomarker-guided treatment claims.

## 6. Conclusion

Cross-cohort evaluation of multi-omics prognostic models should distinguish whether molecular features can be represented, whether a fitted source model transports, whether the same feature set remains informative after local redevelopment, and whether a feature-selection procedure reproduces useful structure in the target cohort.

In TCGA-BRCA and METABRIC, literal transport of the historical RNA and copy-number specifications provided little or negative incremental overall-survival discrimination relative to the transported clinical model. However, retaining the same molecular features while re-estimating model parameters within METABRIC recovered positive incremental discrimination. Random-panel benchmarking showed that this recovery was not uniformly specific to the exact historical feature composition: alternative RNA panels frequently performed similarly, whereas the historical copy-number panel was more strongly ranked within its recovered candidate space.

Independent reconstruction of the dependency-aware selection framework produced a different result. Stable or recurrent molecular structure did not consistently translate into improved overall-survival prediction beyond matched clinical models. A positive RNA contrast was observed for recurrence-free survival, but this secondary endpoint-specific finding requires independent replication.

Together, these results show that failure of a transported molecular model should not automatically be interpreted as failure of its underlying feature set, and that successful local redevelopment should not automatically be attributed to the uniqueness of the selected features. Separating model transport, feature-set information, selection reproducibility, and incremental clinical performance provides a more informative assessment of cross-cohort multi-omics prognostic models.

## Ethics statement

This study used publicly available, de-identified data from TCGA-BRCA and METABRIC. The original cohort publications describe their respective ethics approvals and informed-consent procedures. No new participant recruitment, intervention, or re-identification was conducted. The secondary analyses were performed in accordance with the Declaration of Helsinki and the applicable data-use conditions of the source repositories.

## CRediT authorship contribution statement

**Elena Krikun:** Conceptualization, Methodology, Software, Validation, Formal analysis, Investigation, Data curation, Visualization, Writing – original draft, Writing – review and editing.

**Abedalrhman Alkhateeb:** Conceptualization, Methodology, Supervision, Writing – review and editing.

## Data availability

This study used publicly available, de-identified data from TCGA-BRCA and METABRIC. The source datasets are available through the repositories and access routes described in the corresponding cohort publications. Raw patient-level and large molecular data are not redistributed with this article. Data-source identifiers, processing rules, and scripts for reconstructing the analytic datasets are provided through the project repository, subject to the access conditions of the original data providers.

## Code availability

The analysis code and configuration files for this project are maintained in a public GitHub repository. Raw TCGA-BRCA and METABRIC patient-level data are not included.

## Declaration of generative AI and AI-assisted technologies in the manuscript preparation process

During the preparation of this work, the authors used OpenAI ChatGPT and Anthropic Claude to assist with code development, manuscript organization, and language editing. The authors reviewed and edited all outputs, executed and validated all analysis code locally, and take full responsibility for the content of the article.

## References

[1] M. J. van de Vijver, Y. D. He, L. J. van’t Veer, H. Dai, A. A. M. Hart, D. W. Voskuil, G. J. Schreiber, J. L. Peterse, C. Roberts, M. J. Marton, et al., A gene-expression signature as a predictor of survival in breast cancer, The New England Journal of Medicine 347 (25) (2002) 1999–2009. doi:10.1056/NEJMoa021967.

[2] D. Venet, J. E. Dumont, V. Detours, Most random gene expression signatures are significantly associated with breast cancer outcome, PLoS Computational Biology 7 (10) (2011) e1002240. doi:10.1371/journal.pcbi.1002240.

[3] The Cancer Genome Atlas Network, Comprehensive molecular portraits of human breast tumours, Nature 490 (7418) (2012) 61–70. doi:10.1038/nature11412.

[4] C. Curtis, S. P. Shah, S.-F. Chin, G. Turashvili, O. M. Rueda, M. J. Dunning, D. Speed, A. G. Lynch, S. Samarajiwa, Y. Yuan, et al., The genomic and transcriptomic architecture of 2,000 breast tumours reveals novel subgroups, Nature 486 (7403) (2012) 346–352. doi:10.1038/nature10983.

[5] B. Pereira, S.-F. Chin, O. M. Rueda, H.-K. M. Vollan, E. Provenzano, H. A. Bardwell, M. Pugh, L. Jones, R. Russell, S.-J. Sammut, et al., The somatic mutation profiles of 2,433 breast cancers refine their genomic and transcriptomic landscapes, Nature Communications 7 (2016) 11479. doi:10.1038/ncomms11479.

[6] M. Herrmann, P. Probst, R. Hornung, V. Jurinovic, A.-L. Boulesteix, Large-scale benchmark study of survival prediction methods using multiomics data, Briefings in Bioinformatics 22 (3) (2021) bbaa167. doi: 10.1093/bib/bbaa167.

[7] D. Wissel, N. Janakarajan, A. Grover, E. Toniato, M. Rodríguez Martínez, V. Boeva, SurvBoard: Standardized benchmarking for multi-omics cancer survival models, Briefings in Bioinformatics 26 (5) (2025) bbaf521. doi: 10.1093/bib/bbaf521.

[8] E. Krikun, A. Alkhateeb, A markov blanket-based framework for dependency-aware feature selection in multimodal breast cancer data, Network Modeling Analysis in Health Informatics and Bioinformatics 15 (1) (2026) 158. doi:10.1007/s13721-026-00832-1.

[9] C. Ambroise, G. J. McLachlan, Selection bias in gene extraction on the basis of microarray gene-expression data, Proceedings of the National Academy of Sciences 99 (10) (2002) 6562–6566. doi:10.1073/pnas.102102699.

[10] S. Varma, R. Simon, Bias in error estimation when using cross-validation for model selection, BMC Bioinformatics 7 (2006) 91. doi:10.1186/1471-2105-7-91.

[11] J. L. Haybittle, R. W. Blamey, C. W. Elston, J. Johnson, P. J. Doyle, F. C. Campbell, R. I. Nicholson, K. Griffiths, A prognostic index in primary breast cancer, British Journal of Cancer 45 (3) (1982) 361–366. doi:10.1038/bjc.1982.62.

[12] P. Royston, D. G. Altman, External validation of a cox prognostic model: principles and methods, BMC Medical Research Methodology 13 (2013) 33. doi:10.1186/1471-2288-13-33.

[13] I. Tsamardinos, C. F. Aliferis, A. R. Statnikov, Algorithms for large scale markov blanket discovery, in: Proceedings of the Sixteenth International Florida Artificial Intelligence Research Society Conference, AAAI Press, 2003, pp. 376–381.

[14] F. E. Harrell, K. L. Lee, D. B. Mark, Multivariable prognostic models: issues in developing models, evaluating assumptions and adequacy, and measuring and reducing errors, Statistics in Medicine 15 (4) (1996) 361–387. doi:10.1002/(SICI)1097-0258(19960229)15:4<361::AID-SIM168>3.0.CO;2-4.

[15] P. J. Heagerty, T. Lumley, M. S. Pepe, Time-dependent roc curves for censored survival data and a diagnostic marker, Biometrics 56 (2) (2000) 337–344. doi:10.1111/j.0006-341X.2000.00337.x.

[16] H. Uno, T. Cai, M. J. Pencina, R. B. D’Agostino, L. J. Wei, On the c-statistics for evaluating overall adequacy of risk prediction procedures with censored survival data, Statistics in Medicine 30 (10) (2011) 1105–1117. doi:10.1002/sim.4154.

[17] E. Graf, C. Schmoor, W. Sauerbrei, M. Schumacher, Assessment and comparison of prognostic classification schemes for survival data, Statistics in Medicine 18 (17-18) (1999) 2529–2545. doi:10.1002/(SICI)1097-0258(19990915/30)18:17/18<2529::AID-SIM274>3.0.CO;2-5.

[18] E. W. Steyerberg, A. J. Vickers, N. R. Cook, T. Gerds, M. Gonen, N. Obuchowski, M. J. Pencina, M. W. Kattan, Assessing the performance of prediction models: a framework for traditional and novel measures, Epidemiology 21 (1) (2010) 128–138. doi:10.1097/EDE.0b013e3181c30fb2.

[19] B. Van Calster, D. J. McLernon, M. van Smeden, L. Wynants, E. W. Steyerberg, Calibration: the achilles heel of predictive analytics, BMC Medicine 17 (2019) 230. doi:10.1186/s12916-019-1466-7.

[20] D. G. Altman, P. Royston, What do we mean by validating a prognostic model?, Statistics in Medicine 19 (4) (2000) 453–473. doi:10.1002/(SICI)1097-0258(20000229)19:4<453::AID-SIM350>3.0.CO;2-5.

[21] R. D. Riley, L. Archer, K. I. E. Snell, J. Ensor, P. Dhiman, G. P. Martin, L. J. Bonnett, G. S. Collins, Evaluation of clinical prediction models (part 2): how to undertake an external validation study, BMJ 384 (2024) e074820. doi:10.1136/bmj-2023-074820.

[22] G. S. Collins, K. G. M. Moons, P. Dhiman, R. D. Riley, A. L. Beam, B. Van Calster, et al., Tripod+ai statement: updated guidance for reporting clinical prediction models that use regression or machine learning methods, BMJ 385 (2024) e078378. doi:10.1136/bmj-2023-078378.

