## Supplemental files for "External Evaluation of Multi-Omics Prognostic Models Across TCGA-BRCA and METABRIC: Transportability, Stability, and Incremental Performance"

Table S1: Track B recurrence-free-survival sensitivity analysis. Clinical-only and clinical-plus-omics models were evaluated on the same endpoint- and modality-specific patients. Incremental contrasts and 95% intervals were calculated from unrounded values using 2,000 paired patient-bootstrap samples of the locked repeated out-of-fold predictions. Intervals are conditional on the fitted repeated models and are not full-pipeline bootstrap intervals.

| **Analysis** | $n$ | **Events** | **Clinical** $C$ | **Clinical + omics** $C$ | $\Delta C$ **(95% interval)** | **Interpretation** |
|---|---|---|---|---|---|---|
| RNA | 1,979 | 803 | 0.6496 | 0.6642 | 0.0146 [0.0035, 0.0258] | Positive sensitivity contrast |
| Copy number | 2,078 | 832 | 0.6488 | 0.6547 | 0.0059 [−0.0026, 0.0144] | No reliable improvement |
| Mutation | 2,304 | 932 | 0.6472 | 0.6433 | −0.0040 [−0.0107, 0.0030] | No reliable improvement |
| Methylation | 1,418 | 580 | 0.6507 | 0.6337 | −0.0170 [−0.0319, −0.0017] | Lower than clinical only |
| Reconstructed multimodal | 1,903 | 771 | 0.6505 | 0.6533 | 0.0029 [−0.0136, 0.0204] | No reliable improvement |
